# A multi-dataset evaluation of frame censoring for motion correction in task-based fMRI

**DOI:** 10.1101/2021.10.12.464075

**Authors:** Michael S. Jones, Zhenchen Zhu, Aahana Bajracharya, Austin Luor, Jonathan E. Peelle

## Abstract

Subject motion during fMRI can affect our ability to accurately measure signals of interest. In recent years, frame censoring—that is, statistically excluding motion-contaminated data within the general linear model using nuisance regressors—has appeared in several task-based fMRI studies as a mitigation strategy. However, there have been few systematic investigations quantifying its efficacy. In the present study, we compared the performance of frame censoring to several other common motion correction approaches for task-based fMRI using open data and reproducible workflows. We analyzed eight publicly-available datasets representing eleven distinct tasks in child, adolescent, and adult participants. Performance was quantified using maximum t-values in group analyses, and ROI-based mean activation and split-half reliability in single subjects. We compared frame censoring across several thresholds to the use of 6 and 24 canonical motion regressors, wavelet despiking, robust weighted least squares, and untrained ICA-based denoising, for a total of 240 separate analyses. Thresholds used to identify censored frames were based on both motion estimates (FD) and image intensity changes (DVARS). Relative to standard motion regressors, we found consistent improvements for modest amounts of frame censoring (e.g., 1–2% data loss), although these gains were frequently comparable to what could be achieved using other techniques. Importantly, no single approach consistently outperformed the others across all datasets and tasks. These findings suggest that the choice of a motion mitigation strategy depends on both the dataset and the outcome metric of interest.

## Introduction

High-quality neuroimaging analysis depends in part on minimizing artifacts. Although advancements in hardware and pulse sequence design have reduced many types of noise inherent to functional MRI, other sources remain (Bianciardi et al. 2009). One prominent challenge is artifacts caused by subject head motion. Among other effects, head motion changes the part of the brain sampled by a particular voxel and can introduce changes in signal intensity through interactions with the magnetic field, which add noise to the data and make it harder to identify signals of interest.

The effects of head motion have received recent scrutiny in the context of resting state functional connectivity. Because motion-related artifacts occur in many voxels simultaneously, they can introduce correlations in fMRI time series that are unrelated to BOLD activity, leading to inaccurate estimates of functional connectivity (Power et al. 2015; Satterthwaite et al. 2019). However, spurious activation is also of concern in task-based functional neuroimaging, where it can lead to both false positives, or a lower signal-to-noise ratio that can make it harder to detect a true activation of interest. As such, motion in task-based fMRI potentially introduces a combination of both Type I and Type II errors.

Rigid body realignment—a mainstay of fMRI analysis for decades—goes some way towards improving correspondence across images (Ashburner and Friston 2004), but does not remove extraneous signal components introduced by movement (Friston et al. 1996). A common approach for mitigating motion-related artifacts is to include the 6 realignment parameters (translation and rotation around the X, Y, and Z axes, reflecting estimated participant motion) as nuisance regressors in first-level models.

Beyond motion parameter inclusion, several data-driven strategies have been developed to reduce the influence of high-motion scans on estimated activations. Wavelet decomposition identifies artifacts by exploiting their non-stationarity across different temporal scales (Patel et al. 2014). The method has been applied in resting state studies but is also applicable to task-based data. Independent component analysis (Pruim et al. 2015) identifies artifacts based on the spatial distribution of shared variance. In robust weighted least squares (Diedrichsen and Shadmehr 2005), a two-pass modeling procedure is used to produce a collection of nuisance regressors which are then included in the final analysis to weight frames by the inverse of their variance (that is, downweighting frames with high error).

An alternative motion correction strategy is “scrubbing” or “frame censoring” (Lemieux et al. 2007; Siegel et al. 2014). In this approach, bad scans are identified and excluded from statistical analysis. One approach is to do so by modeling them in the general linear model using nuisance regressors (i.e., “scan-nulling regressors” or “one-hot encoding”). Although frame censoring has received considerable interest in resting state fMRI over the past several years (Power et al. 2012; Gratton et al. 2020a), it has not seen widespread use in the task-based fMRI literature. Censoring approaches involve some effective data loss, in that censored frames do not contribute to the task-related parameter estimates, and that columns introduced to the design matrix to perform censoring reduce the available degrees of freedom. There are different ways to quantify “bad” scans, and choosing both an appropriate metric and associated threshold can also be challenging. Thus, additional information over what threshold should be used for identifying bad frames—and relatedly, how much data is lost vs. retained—is necessary to make informed decisions.

Although several published studies compare differing correction strategies (Ardekani et al. 2001; Oakes et al. 2005; Johnstone et al. 2006), a drawback of prior work is that evaluation was often limited to a single dataset (see **Supplemental Table 1**). The degree to which an optimal strategy for one dataset generalizes to other acquisition schemes, tasks, or populations is not clear. With the increased public availability of neuroimaging datasets (Poldrack et al. 2013; Markiewicz et al. 2021), the possibility of evaluating motion correction approaches across a range of data has become more feasible.

In the present work, we sought to compare the performance of identical pipelines on a diverse selection of tasks, using data from different sites, scanners, and participant populations. Although our primary interest was frame censoring, we considered seven different motion-correction approaches:

1. six canonical head motion (i.e., “realignment parameter”) estimates (RP6)
2. 24-term expansions of head motion estimates (RP24)
3. wavelet despiking (WDS)
4. robust weighted least squares (rWLS)
5. untrained independent component analysis (untrained ICA; uICA)
6. frame censoring based on frame displacement (FD)
7. frame censoring based on variance differentiation (DVARS)

This list is not exhaustive but representative of approaches that are currently used and feasible to include in an automated processing pipeline.

Because it is impossible to determine a “ground truth” result with which to compare the effectiveness of these approaches, we instead considered four complementary outcome metrics: 1) the maximum group t-statistic both across the whole-brain and in a region-of-interest (ROI) relevant to the task; 2) the average parameter estimates from within the same ROI; 3) the degree of test-retest consistency exhibited by subject-level parametric maps; and 4) the spatial overlap of thresholded group-level statistical maps. These metrics are simple to define yet functionally meaningful, and can be applied to data from almost any fMRI study. In our view, Dice quantifies replicability, the mean ROI value quantifies effect size (signal), and maximum-t quantifies signal-to-noise (effect size penalized by variance).

## Methods

### Datasets

We analyzed eight studies obtained from OpenNeuro (Markiewicz et al. 2021), several of which included multiple tasks or multiple participant groups. As such, the eight selected studies provided a total of 15 datasets. The selection process was informal, but studies given priority included: 1) a clearly-defined task; 2) a sufficient number of subjects to allow second-level modeling; 3) sufficient data to make test-retest evaluation possible; and 4) a publication associated with the data describing a result to which we could compare our own analysis.

A summary of the eight datasets selected is shown in **Table 1** (acquisition details are provided in **Supplemental Table 2**). Additional information, including task details, modeling/contrast descriptions compiled from publication(s) associated with a given study, and any data irregularities encountered during analysis, is provided in the **Supplemental Materials**.

**Table 1.** Summary of datasets analyzed.

| Dataset | Reference | Task and design | # subs | Age range | FD (median ± SD) | frames per subject |
| --- | --- | --- | --- | --- | --- | --- |
| ds000102 | Kelly et al. (2008) | flanker (E) | 22 | 22-50 | 0.11 ± 0.12 | 284 |
| ds000107 | Duncan et al. (2009) | 1-back (B) | 43 | 19-38 | 0.08 ± 0.14 | 323 |
| ds000114 | Gorgolewski et al. (2013a) | motor (B) | 10 | 50-58 | 0.14 ± 0.16 | 360 |
| | | covert verb (B) | 10 | 50-58 | 0.11 ± 0.11 | 338 |
| | | overt word (B) | 10 | 50-58 | 0.13 ± 0.12 | 144 |
| | | line bisection (B) | 9 | 50-58 | 0.13 ± 0.18 | 468 |
| ds000228 | Richardson et al. (2018) | film viewing (E) | 122 | 3.5-12 | 0.21 ± 0.93 | 164 |
| | | | 33 | 18-39 | 0.18 ± 0.27 | 164 |
| ds001497 | Lewis-Peacock and Postle (2008) | face perception (E) | 10 | 19-32 | 0.11 ± 0.12 | 1146 |
| ds001534 | Courtney et al. (2018) | food images (E) | 42 | 18-22 | 0.10 ± 0.16 | 552 |
| ds001748 | Fynes-Clinton et al. (2019) | memory retrieval (E) | 21 | 10-12 | 0.16 ± 0.36 | 438 |
| | | | 20 | 14-16 | 0.12 ± 0.17 | 438 |
| | | | 21 | 20-35 | 0.08 ± 0.17 | 438 |
| ds002382 | Rogers et al. (2020) | word recognition (E) | 29 | 19-30 | 0.14 ± 0.35 | 710 |
| | | | 32 | 65-81 | 0.30 ± 0.34 | 710 |

B = blocked design, E = event-related design

### Analysis

All scripts used in the study are available at https://osf.io/n5v3w/. Analysis was performed using Automatic Analysis version 5.4.0 (Cusack et al. 2015; RRID: SCR_003560), which scripted a combination of SPM12 (Wellcome Trust Centre for Neuroimaging) version 7487 (RRID: SCR_007037) and the FMRIB Software Library (FSL; FMRIB Analysis Group) (Jenkinson et al. 2012) version 6.0.1 (RRID: SCR_002823). BrainWavelet Toolbox v2.0 (Patel et al. 2014) was used for wavelet despiking and rWLS version 4.0 (Diedrichsen and Shadmehr 2005) for robust weighted least squares.

To the extent possible, we used the same preprocessing pipeline for all datasets (**Figure 1a**). Briefly, structural and functional images were translated to the center of the scanned volume and the first four frames of each session were removed in functional images to allow for signal stabilization. This was followed by bias correction of the structural image, realignment, coregistration of the functional and structural images, normalization into MNI space using a unified segmentation approach (Ashburner and Friston 2005) resampled to 2 mm isotropic voxels, and smoothing of the functional images using an 8 mm FWHM Gaussian kernel.

**Figure 1.**
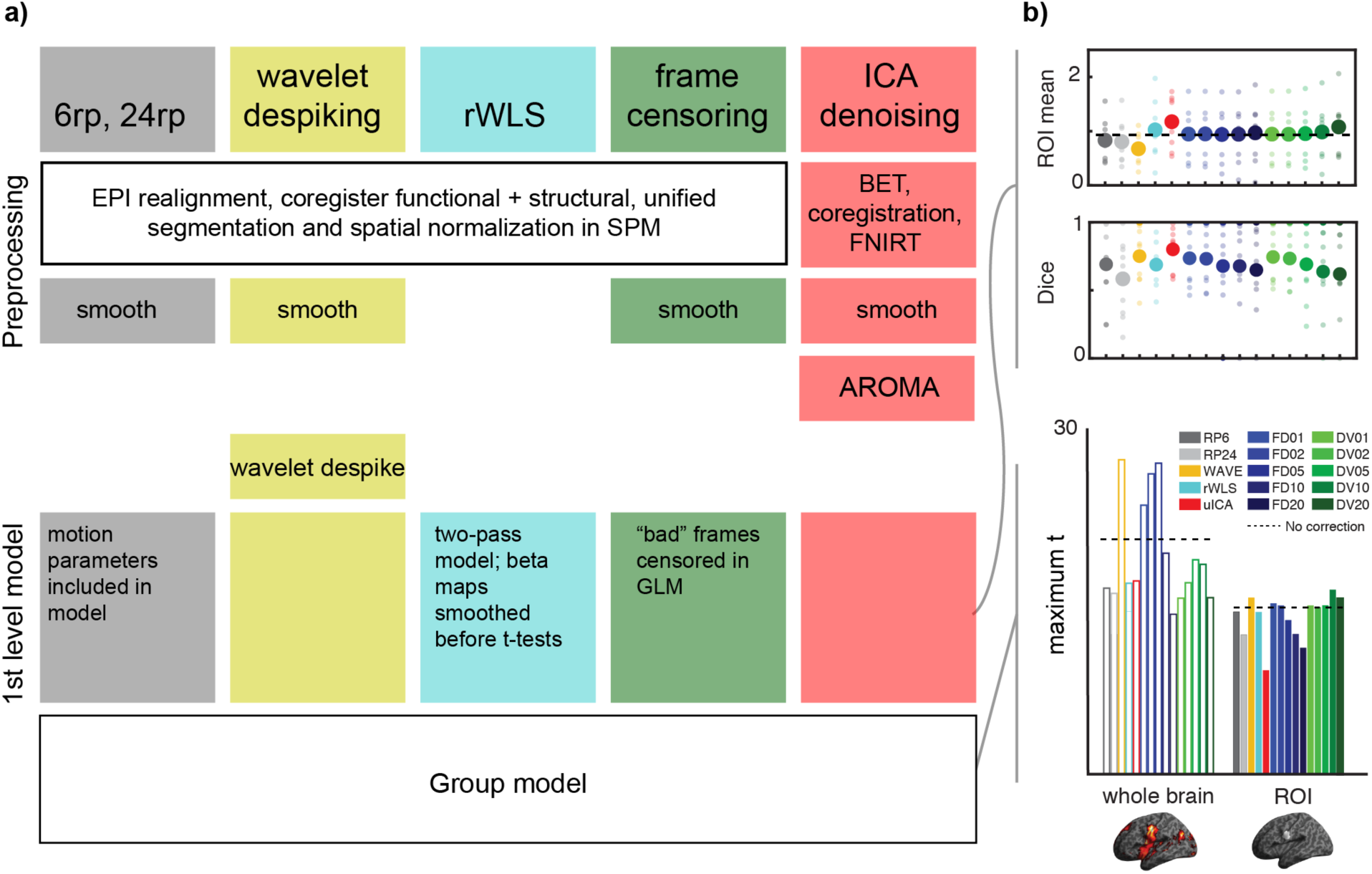
Schematic of processing pipeline and outcome measures. **(a)** Summary of preprocessing and model steps in common and differing across motion correction strategies. **(b)** Following statistical modeling, outcomes are summarized in mean parameter estimates and Dice overlap of thresholded single-subject maps (top) and maximum t-value from the group analysis (bottom). Dashed lines represent values obtained without motion correction.

Functional images were corrected for motion artifacts using each of the following approaches: 1) inclusion of six canonical motion estimates in the first-level model as nuisance regressors; 2) inclusion of 24 nuisance regressors based on a second-order expansion of the motion estimates and first derivatives; 3) wavelet despiking, 4) robust weighted least squares; 5) untrained ICA denoising; 6) frame censoring based on framewise displacement (FD); or 7) differential variance (DVARS) thresholding (FD/DVARS thresholding is described below).

Statistical modeling was performed in SPM for all motion correction approaches. First-level modeling included a contrast of interest described in a publication associated with the dataset for evaluation, followed by second-level analysis to produce group-level statistical maps. All first- and second-level t-maps were thresholded at a voxelwise threshold of p < 0.001 (uncorrected).

Minor pipeline modifications were required for robust weighted least squares, wavelet despiking, and untrained ICA denoising. As recommended by developers of the rWLS toolbox, unsmoothed data was used for variance estimation and contrast maps were smoothed after modeling. For wavelet despiking, functional images were rescaled to a whole-brain median of 1000 across all frames prior to processing. The default toolbox settings (wavelet: d4, threshold: 10, boundary: reflection, chain search: moderate, scale number: liberal) were used. Finally, untrained ICA-based denoising was implemented using ICA-AROMA (Pruim et al. 2015) with additional processing steps performed within FSL. Briefly, the unsmoothed coregistered functional image was demeaned, detrended, smoothed, and then nonlinearly warped to the FSL 2 mm MNI152 template using FNIRT. The normalized functional image was then passed to AROMA for denoising. This ICA implementation is not based on training data and so we refer to it as “untrained” ICA to distinguish it from other ICA-based denoising approaches.

### Evaluation of Motion Correction Performance

Three measures were used to quantify the performance of each motion correction strategy, illustrated in **Figure 1b**: 1) maximum t-value, 2) effect size, and 3) subject replicability. In the first measure, the maximum t-value occurring in the group level parametric map was extracted both at the whole-brain level and also within a region-of-interest relevant to the task. The effect size was quantified as the mean of all voxels within the ROI for each subject using the first-level beta maps. To evaluate subject replicability, multisession data were treated as a test-retest paradigm (the first session statistical map was compared to the second session in studies having fewer than three sessions; even-numbered versus odd-numbered sessions were compared otherwise). Replicability was quantified as the Dice coefficient of thresholded first-level t-maps (p< 0.001, uncorrected) in each subject (restricted to the ROI).

### FD and DVARS Thresholding

Motion correction approaches based on frame censoring required quantification of motion artifacts which could then be subjected to thresholding. Both framewise displacement (FD) and differential variance (DVARS) were used. Framewise displacement was calculated as the sum of the six head motion estimates obtained from realignment, with a dimensional conversion of the three rotations assuming the head is a 50 mm sphere (Power et al. 2012). DVARS was calculated as the root-mean-squared of the time difference in the BOLD signal calculated across the entire brain (Smyser et al. 2011). As shown in **Figure 2a**, both metrics closely tracked artifacts apparent in voxel intensities and also each other. Although FD and DVARS in a given session tended to be correlated (**Figure 2b**), they were not identical and could exhibit slightly different time courses and relative peak amplitudes (**Supplemental Figure S1**). As such, we explored the use of both measures.

**Figure 2.**
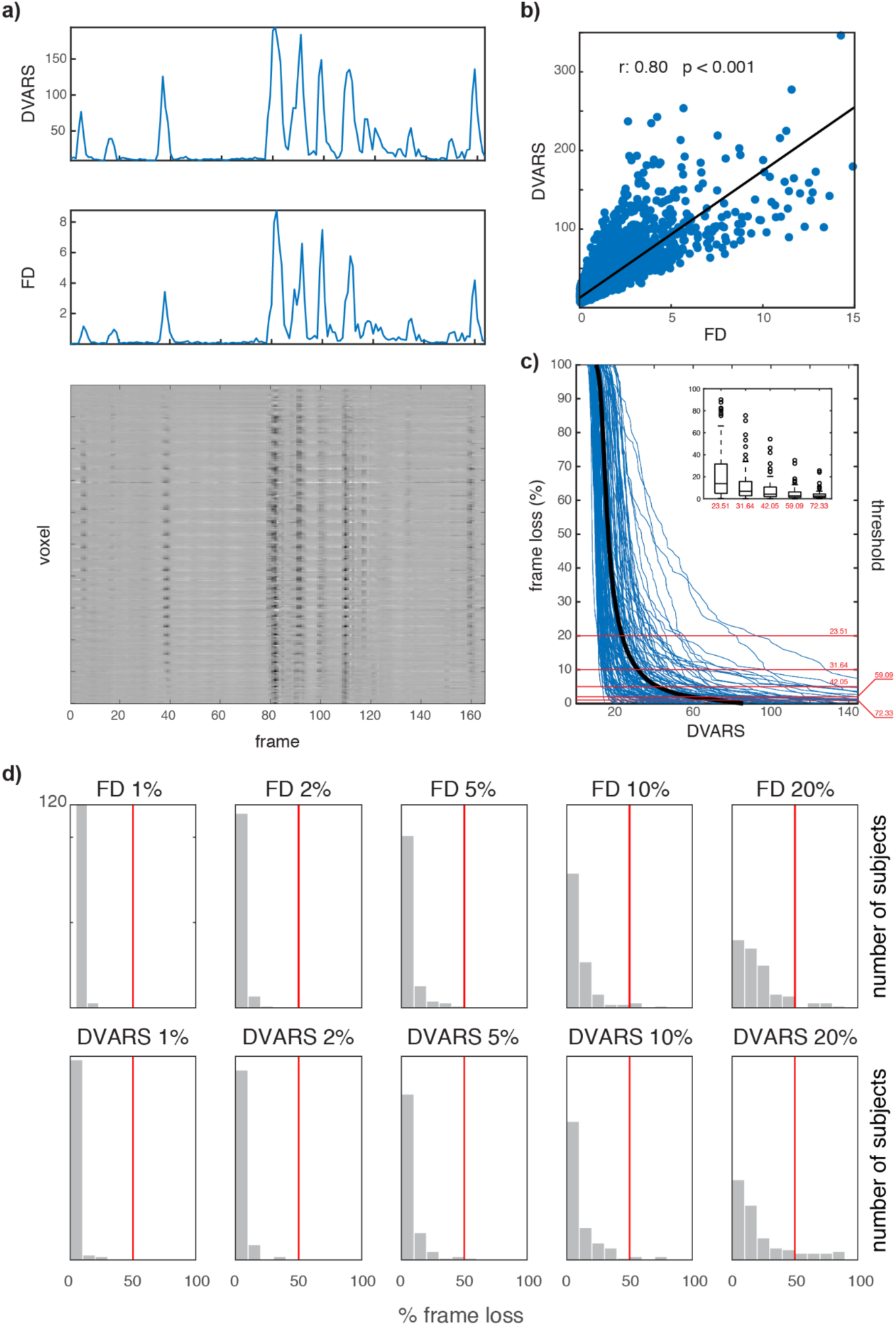
Calculation of censoring thresholds in an example dataset (ds000228, children). **(a)** Representative grayplot (*bottom;* Power 2017) showing 500 randomly selected gray matter voxels. DVARS and FD for this session are plotted above. Spikes in the metrics identify frames contaminated by artifacts. **(b)** DVARS and FD are correlated but exhibit differing amplitudes and time courses. As such, the use of both measures was explored. **(c)** Metric values (here shown for DVARS) used for censoring were determined by plotting frameloss for each subject as a function of threshold (thin blue traces). Interpolation of the mean response (thick black trace) gives metric values corresponding to a data loss of 1%, 2%, 5%, 10%, or 20%. Box plot (*inset*) summarizes the results across all subjects at each threshold (box: 25-75% percentiles; crosses: >3 SD outliers). **(d)** Histograms of data loss at each FD or DVARS target value across all subjects in this dataset. Red line indicates 50% frame loss.

Thresholds were determined by calculating FD and DVARS across all sessions in all subjects, which allowed values to be identified that resulted in 1%, 2%, 5%, 10%, and 20% frame violations across the entire dataset (**Figure 2c**). We adopted this strategy rather than using a fixed value of FD or DVARS for several reasons. First, FD and DVARS magnitudes change with the TR of the data, because the TR is the sampling rate (for a given movement, sampling more rapidly will give smaller FD values, even though the total motion is the same). Secondly, different calculations of FD provide different values (Jenkinson et al. 2002; Power et al. 2012; Van Dijk et al. 2012), and thus any absolute threshold would necessarily be metric-specific. Finally, datasets differ in their tasks and populations, and we anticipated that a single threshold would not be suitable for all datasets. We, therefore, employed the frame-percent thresholding strategy to obtain a reasonable range of results in all studies examined. To be clear, we do not propose a fixed-percent data loss approach as a “production” strategy. Indeed, a conventional denoising approach would be to select an absolute threshold of acceptable motion based on experience (or preliminary data) from the target subject pool, task, scanner, and so on. However, given the variety of datasets examined here, we had no *a priori* guide as to what threshold values to use: Any fixed selection might censor no frames in some studies and too many in others. Therefore, we employed the frame-percent thresholding strategy to obtain an informative range of results in all datasets. The threshold values that resulted from percent data loss targeting in these datasets are shown in **Supplemental Figure S2** and listed in **Supplemental Table 3**. The amount of data censored in each participant in a single study is shown in **Figure 2d**, and for all studies in **Supplemental Figure S3**.

To implement frame censoring, first-level modeling was repeated for each threshold with a separate delta function (i.e., a scan-nulling regressor) included in the design matrix at the location of each violation, which effectively removes the contribution of the targeted frame from the analysis. Although some prior studies of motion correction have censored one or more frames prior to or following threshold violations (e.g., “augmentation” of Siegel et al. 2014), we did not pursue such variations to avoid further expanding what was already a rather large parameter space.

### Region of Interest Definition

A task-relevant ROI for each study/task was defined in one of three ways: 1) a 5-mm sphere (or spheres) centered at coordinates reported in a publication associated with the dataset; 2) a whole-brain Z-mask generated by a task-relevant search term (e.g., “incongruent task”) in NeuroQuery (Dockès et al. 2020) and thresholded Z > 3; or 3) a binarized probability map in the SPM Anatomy Toolbox (Eickhoff et al. 2005) for a task-relevant brain structure or anatomical region (e.g., “V2”). Additional details on the ROI definition used in each analysis are provided in the **Supplemental Materials**.

## Results

Performance of the motion correction strategies organized by dataset is shown in **Figure 3**. Each panel includes a second-level thresholded t-map at the upper left (p < 0.001, uncorrected) using the “RP6” approach (six canonical motion parameters included as nuisance regressors). A contrast descriptor is given below the map. The ROI used for evaluation is shown at lower left with the source listed under the rendered image.

**Figure 3.**
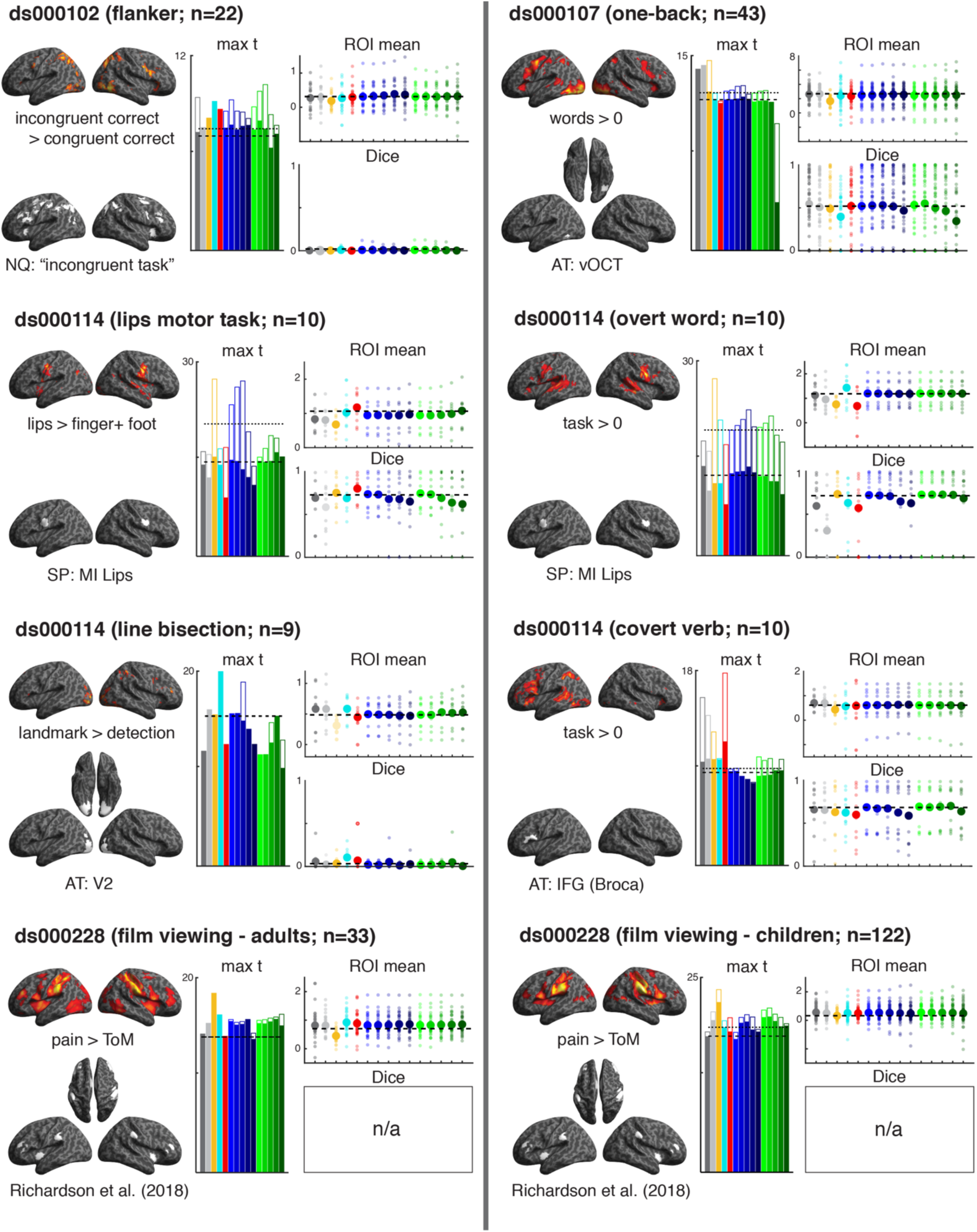
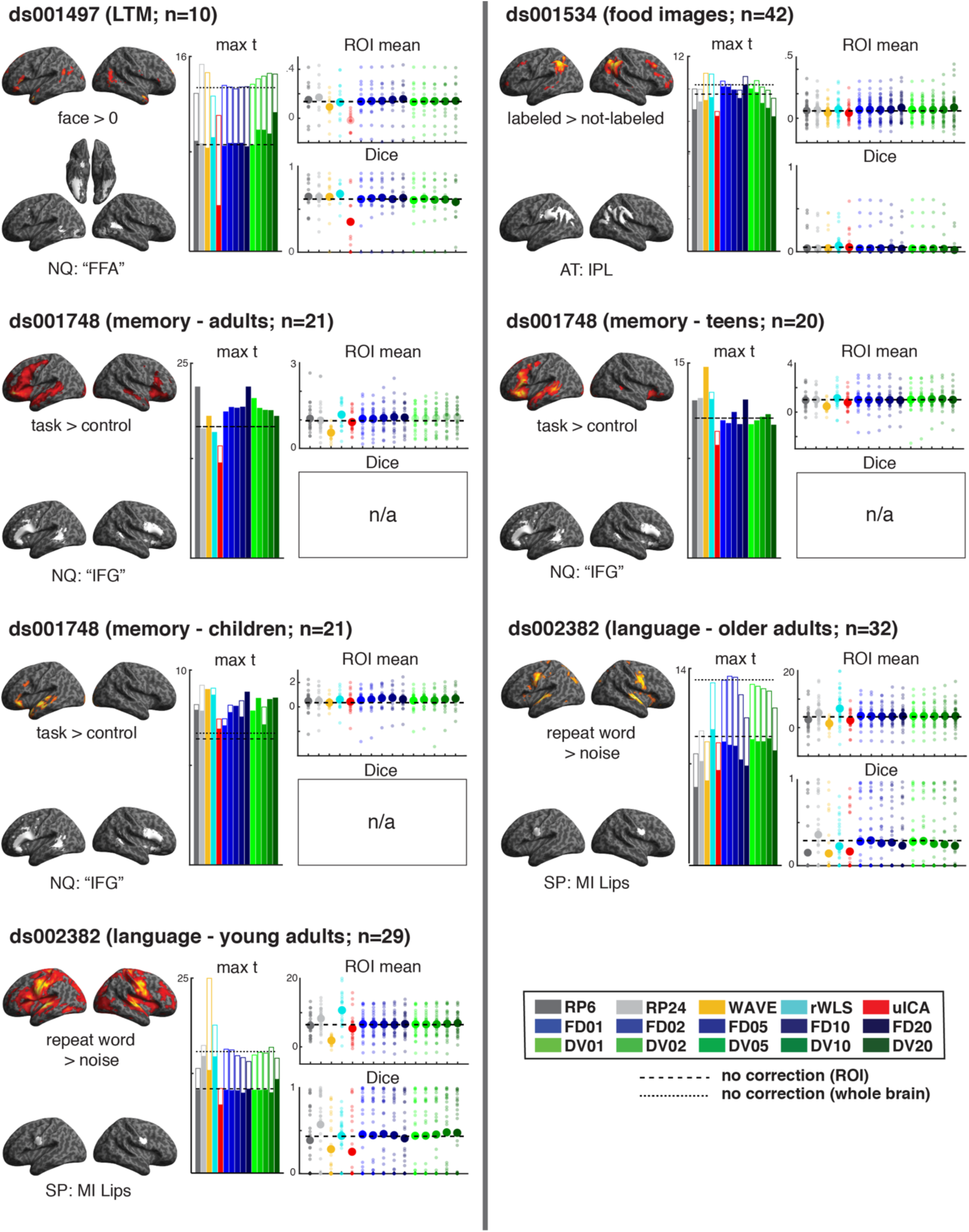
Summary of motion correction algorithm performance for all datasets examined in the study. Each panel includes a representative thresholded group t-map at left (p=0.001, uncorrected) for the given contrast with the ROI used for evaluation plotted below (AT: Anatomy Toolbox probability map; NQ: NeuroQuery search term; SP: 5 mm sphere centered at the described landmark. The ROI used in the analysis of ds000228 is defined in Table 2 of Richardson et al. (2018)). At the center, ROI-restricted maximum t-values are superimposed on whole-brain results for each motion correction approach. Plots at right show individual-subject mean ROI effect size (*top*) and Dice coefficient for a split-half test-retest evaluation (*bottom*). Datasets that did not permit test-retest evaluation are noted “n/a.” Horizontal dashed reference lines indicate the value obtained when no motion correction was used (dashed: ROI-constrained; dotted: whole brain).

These results show there is substantial variability in motion correction approaches, with performance depending both on the data under consideration and the chosen performance metric. However, some general trends are apparent. Wavelet despiking tends to offer the best maximum t-value in both the whole-brain and ROI-constrained evaluation, with robust weighted least squares also exhibiting good performance (note the ROI-constrained maximum t-value, shown in filled bars, are superimposed on the whole-brain results, shown in open bars in **Figure 3**). Conversely, untrained ICA gives consistently poorer results although it offers the best maximum t-value in the ds000114 covert verb task. Performance of FD and DVARS frame censoring was highly variable, with the application of increasingly stringent thresholds improving performance in some datasets while decreasing it in others. A somewhat consistent result is a loss of performance at the highest (20%) FD or DVARS threshold. As a rule, frame censoring performed better than RP6 and RP24 motion correction, although RP6 is competitive (if not optimal) in both ds000107 and ds001748.

The mean effect size shown in these results was largely insensitive to the selected motion correction approach. The two exceptions are wavelet despiking and untrained ICA, which produce consistently smaller values than the other approaches. This may reflect suboptimal parameter selection in these algorithms (see **Discussion**). Robust weighted least squares offers competitive results in all datasets and notably superior results in ds002382 and the ds000114 overt word task. FD and DVARS frame censoring neither improved nor degraded results regardless of threshold, producing a mean effect size indistinguishable from both the RP6 and RP24 approaches save for a few individual subjects.

The test-retest results also demonstrate a great deal of variability. The Dice coefficients exhibit substantial inter-subject differences, resulting in a mean performance that is similar across all motion correction strategies. However, excluding ds000102, ds001534, and the ds000114 line bisection task, all of which unfortunately provided an uninformative test-retest quantification, some trends can be identified. There is a decrease in both the FD and DVARS frame censoring results, especially at 20% thresholding. In general, all differences were minor, save for untrained ICA which performs notably better in the ds000114 motor task and notably worse in ds001487. The reason why three datasets exhibit poor performance in a test-retest paradigm is unclear. Although ds000114 had a relatively small subject pool (n=10), both ds000102 and ds001534 used a larger sample size (n=22 and n=42, respectively). Whatever the cause, it appears to be unrelated to the choice of motion correction, as in these exceptions all strategies performed equally well (or equally poorly, as it were).

A summary of univariate results is shown in **Figure 4a**, in which mean values of all four performance metrics are plotted. Several of the trends noted in the individual datasets remain apparent. For example, wavelet despiking gave the largest whole-brain maximum t-value, whereas robust weighted least squares resulted in the best ROI-constrained performance. Light-to-moderate frame censoring resulted in improvement which then declined as more aggressive thresholding was applied. Robust weighted least squares produced the largest average effect size. Wavelet despiking and untrained ICA produce poor results as measured by this metric. Test-retest performance is generally poorer for most motion correction strategies than that obtained using no motion correction, although rWLS exhibits good performance as measured by this metric.

**Figure 4.**
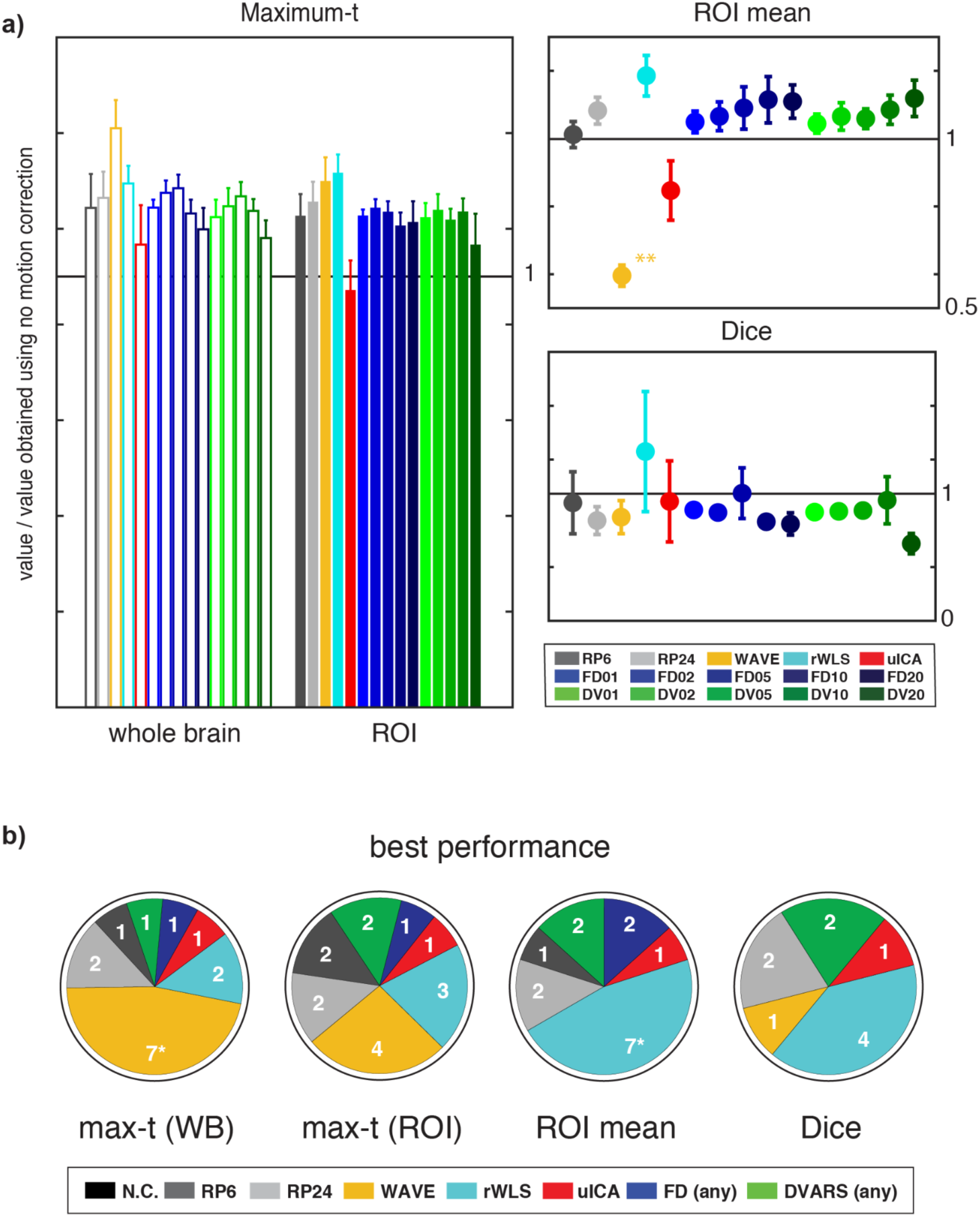
Summary statistics. **(a)** Whole-brain and ROI-restricted maximum t-values (*left*) mean effect size (*upper right)*, and test-retest Dice coefficient (*lower right)* averaged across all datasets. Values were normalized to the no-correction values in the same dataset prior to pooling. **(b)** Performance summarized as a count of the datasets a given approach gave best results measured by the four performance metrics (*p < 0.05).

An omnibus ANOVA identified a significant difference in the maximum-t data, however Scheffe post hoc testing found no significant pairwise differences (p > 0.05). Both omnibus and post hoc testing of the mean ROI effect size show wavelet despiking differed significantly from all other approaches (p < 0.001). No significant differences were found in the test-retest Dice data.

A count summary of best algorithm performance is shown in **Figure 4b**, in which the best performing motion correction approach for each metric was identified in each of the 15 datasets and the resulting proportions plotted as pie charts. The general trends evident in the averaged results are also apparent in these data although some additional features emerge. Robust weighted least squares offered the best performance on many datasets. Wavelet despiking gave the best maximum t-value in approximately half (whole-brain) or one quarter (ROI-constrained) of the studies. Untrained ICA gave best results across all four metrics in at least one dataset. Frame censoring performed similarly using either FD or DVARS. Finally, the performance of the RP6 and RP24 approaches are middling, producing the best maximum t-value on only one or two datasets and, with one exception, never producing the best ROI mean or test-retest results. However, of these results, only the maximum-t performance of wavelet despiking and rWLS ROI mean effect size were statistically significant (p < 0.05).

Given the substantial variability in motion correction results across datasets, we next explored whether there may have been systematic differences between datasets that affected motion correction performance. We first calculated the pairwise similarity of thresholded (voxelwise p < .001) group maps from each dataset using Dice overlap (**Figure 5**). A consistent finding was a generally lower overlap between untrained ICA and the other motion correction approaches. Additionally, RP6, RP24, and rWLS tended to overlap less with other motion correction approaches and more with one another, although exceptions can be noted. Results for most datasets are generally mixed, although ds000228 (adults), ds001748 (adults), and ds002382 (young adults) exhibit high overlap for all motion correction approaches (with the exception of untrained ICA).

**Figure 5.**
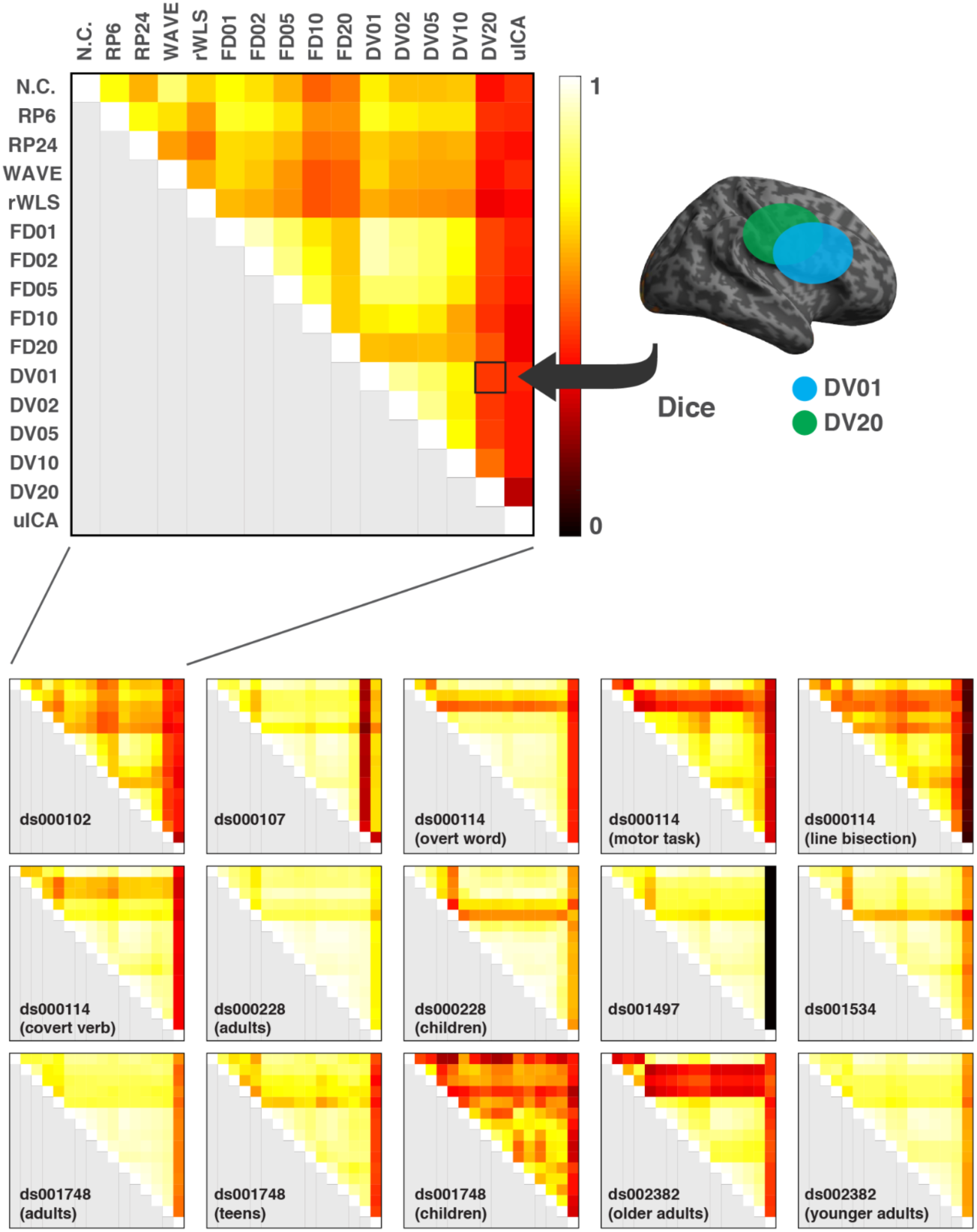
Quantification of overlap in group-level maps. Dice coefficients computed from group-level thresholded statistical maps obtained using each pair of motion correction strategies were assembled into a 15×15 Dice matrix. Overlap shown here for DV01 and DV20 is illustrative.

Having generated Dice overlap maps for each dataset, we then explored the higher-order relationship between datasets using representational similarity analysis (Kriegeskorte et al. 2008). We first calculated the distance between each Dice matrix using Pearson correlation, creating a representational dissimilarity matrix (RDM) based on these distances (**Figure 6a**). We then used multidimensional scaling (MDS) to visualize the relationship between datasets. A plot of the data in the first two eigen-dimensions is shown in **Figure 6b**. Dataset ds000107 appears at the right edge of the space, as might be predicted by a visual review of the RDM. However, the other datasets present no distinct pattern. A plot of the data using the first three dimensions similarly exhibited no distinct features, as did an examination of all 2D projections using the first five eigen-dimensions (see **Supplemental Material Figure S4**).

**Figure 6.**
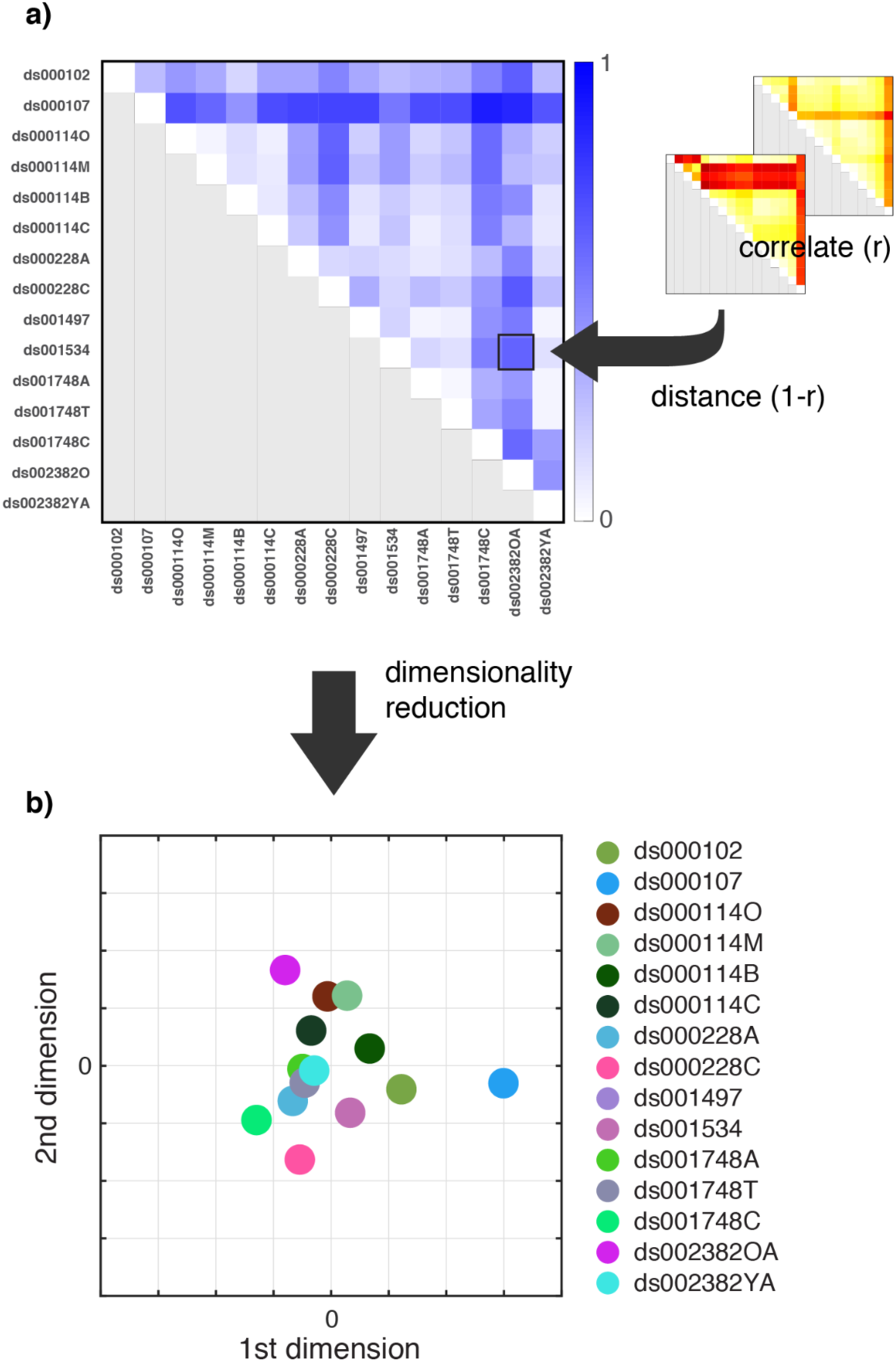
Multivariate analysis of group-level overlap. (a) Representational dissimilarity matrix (RDM) illustrating the distance between motion correction patterns for each of the 15 data sets shown in Figure 5. (b) Using multidimensional scaling (MDS) we visualized the relative distances between data sets in a reduced number of dimensions. Plotting the first two dimensions partially segregates ds000107 (cyan dot) but does not suggest other organization of the datasets. Plots of other low-dimensional projections were qualitatively similar (see also Supplemental Figure S4).

## Discussion

We explored the performance of a variety of approaches to correcting motion-related artifacts in task-based fMRI. The studies examined represent a broad range of task domains, including sensory, motor, language, memory, and other cognitive functions, with participants varying in age, sex, and other characteristics. Although we set out expecting to find converging evidence for an optimal strategy, instead our results demonstrate that the performance of motion correction approaches depends on both the data and the outcome of interest. We review our selected metrics below—whole-brain and ROI-restricted maximum t-value, mean effect size, and test-retest repeatability—followed by some general comments on each motion correction approach.

### Comparing outcome metrics

The use of whole-brain maximum t-value measured in group-level statistical maps has the advantage that it requires few assumptions about the data or the expected pattern of activity. However, we did not observe a consistent pattern regarding which motion correction approach optimized the whole-brain maximum t-value. The disparity was even evident between different participant groups within a given study. For example, wavelet despiking had the highest whole-brain t statistic in ds0001748 in teens but RP6 offered better performance in adults.

In addition to whole-brain statistics, we examined maximum t-values within a selected region of interest. Our rationale for doing so was that researchers interested in task-based effects frequently have prior intuitions about where the most informative results are localized. We found that motion correction approaches can exhibit substantially different whole-brain and ROI-restricted performance. In the ds000114 overt word task, for example, RP6 offered best performance within the motor cortex but poor performance in a whole-brain evaluation.

Furthermore, frame censoring performance improved in some datasets but degraded in others as more stringent thresholding was applied. Obviously, a challenge inherent in such an evaluation is the actual ROI selection. Although we believe our choices are sensible, the selection of a different ROI set may well result in a different overall view of performance.

To complement these group-level measures, we also considered two single-subject metrics: mean effect size and test-retest repeatability. Effect size permits an examination of parameter estimates, and our use of averaging offers a direct and simple quantification. However, with the exceptions of wavelet despiking and aggressive frame censoring (revisited below), we observed that effect size was largely insensitive to the choice of motion correction strategy, although less than the variability observed in the maximum t-value. This suggests the main effect of different motion correction approaches is a differential reduction in model error variance. If parameter estimation is the primary result of interest, then the choice of motion correction strategy may not be critical.

The test-retest evaluation was perhaps the least helpful result, with the performance of all motion correction approaches essentially indistinguishable under this metric. Although the outcome is disappointing, it should be noted that many of the studies included here were not designed to include a split-half repeatability analysis. It may be that more data per subject may be needed for this metric to be informative. In that sense, our analyses speak to the general challenges of obtaining reliable single-subject data in fMRI (Smith et al. 2005; Bennett and Miller 2010; Gorgolewski et al. 2013b; Elliott et al. 2020), at least under conventional scanning protocols (Gratton et al. 2020b).

### Comparing Motion Correction Approaches

No single motion correction approach exhibited optimal performance on all datasets and all metrics. Algorithm performance did not appear to be systematically related to the nature of the task, acquisition parameters, nor any feature of the data that we could identify.

Interestingly, computationally-intensive approaches did not necessarily perform better than basic corrective measures. For some datasets, including six motion estimates as continuous nuisance regressors—a standard approach used in functional imaging for decades—performed as well or better than more sophisticated algorithms that have emerged in recent years. Increasing the head motion estimate from a 6-to a 24-parameter expansion led to an improvement in some data but poorer results in others. Although such results are rather counterintuitive, we can provide a few observations, even if these data do not currently permit conclusive recommendations.

Two motion correction approaches that showed generally strong performance were wavelet despiking (WDS) and robust weighted least squares (rWLS). Together, these approaches offered the best performance in approximately half of the datasets across all performance metrics (**Figure 4b**). In a statistical sense, robust weighted least squares might be seen as an optimal solution in that it uses the error in the model to differentially weight time points, reducing the influence of motion on parameter estimates. However, we found that other motion correction strategies offered similar, or superior, performance in several instances. One reason might be that rWLS linearly weights time points inversely related to their variance. To the degree that motion artifacts include a nonlinear component, linear weighting may not adequately (or not optimally) remove all of the artifacts.

In contrast to the good performance of wavelet despiking as measured by maximum t-value, it gave notably low scores on mean effect size. However, this finding may simply reflect data scaling specific to the toolbox implementation. It should also be noted the wavelet despiking toolbox offers 20 wavelets and additional options that control algorithm behavior such as thresholding and chain search selection. The results obtained here are what can be expected using the default settings recommended by the toolbox developers, which includes a median 1000 rescaling of the functional data (and hence the lower parameter estimate). Thus, numeric comparison to other approaches (that do not include rescaling) is problematic. It also may be possible to improve performance—including obtaining effect sizes concomitant with other motion correction approaches—by tuning the algorithm.

One unexpected result was the relatively poor performance of ICA denoising. Although individual exceptions exist, the approach produced consistently low scores on all evaluation metrics. However, it should be emphasized that we implemented ICA denoising using FSL’s ICA-AROMA, specifically selected because it does not require classifier training. More sophisticated ICA denoising tools such as MELODIC or ICA-FIX involve a visual review of training data to generate a set of noise classifiers based on the temporal, spatial, and frequency characteristics of identified artifacts (Salimi-Khorshidi et al. 2014; Griffanti et al. 2017). These options were not considered presently because we sought to evaluate tools for motion correction that could be implemented within a completely automated pipeline. The potential of ICA, in general, for denoising task-based data should not be dismissed; rather, our results only indicate that the use of *untrained* ICA is probably suboptimal compared to other options, many of which are also less computationally intensive.

#### Frame Censoring

Frame censoring has appeared in several task-based studies (O’Hearn et al. 2016; Bakkour et al. 2017; Davis et al. 2017). In fact, it was an experience with frame censoring in the analysis of in-scanner speech production (Rogers et al. 2020) that motivated our interest in comparing motion correction approaches. We found that modest levels of frame censoring (e.g., 2–5% data loss) revealed a regional activation in high-motion subjects that appeared in low-motion subjects but was not apparent when standard (RP6) motion correction was used. This suggested that using a discrete rather than a continuous nuisance regressor may better preserve task-related variance in some applications. However, a more nuanced picture emerges from the present results, which suggest frame censoring is neither universally superior to nor worse than RP6. One possibility is that frame censoring performance involves a complex interaction between data quantity and data quality. Because each censored frame introduces an additional regressor to the design matrix, eventually the reduction in error variance may be overwhelmed by a loss of model degrees of freedom, or by the effective loss of task-related data. This is anecdotally supported by a decline in many of the metric results observed here at the most stringent FD or DVARS thresholds, an effect that was even more pronounced when 40% maximal censoring was explored in pilot work (data not shown).

One might argue that frame censoring should be based on a selected fixed threshold rather than a targeted percent data loss. The present results offer somewhat mixed support for such a position. We investigated applying a (fixed) FD threshold of 0.9 to these data (**Supplemental Figure S2**), as used by Siegel and colleagues (2014) in their exploration of frame censoring and as well as other studies (e.g., Davis et al. 2017). In most of the datasets considered here, a 0.9 FD threshold would have resulted in less than 1% of frames being censored. This would be a reasonable amount of data loss and might lead to some improvements compared to a standard RP6 approach (although we did not test this directly). However, ds000228 (adults), ds001748 (teens), and ds002382 (YA) would have incurred a 1-2% data loss, ds001748 (child) and ds002382 (OA) approximately 5% data loss, and ds000228/child approximately 13% data loss. These outcomes do not correspond to the best performance obtained across all approaches. Whole-brain or ROI-constrained maximum-t metrics are optimal at these values in some, but not all, datasets. Mean effect size and Dice coefficients add little to the evaluation as they appear largely insensitive to frame censoring thresholds in this range. Taken together, these results suggest there is no single threshold value that will optimize frame censoring for all datasets and outcome measures. Although for individual investigators it may indeed make more sense to develop censoring criteria based on the range of FD or DVARS values present in their specific data, we also suggest that considering the amount of data lost at a chosen threshold is a useful metric to take into consideration.

#### Effects of FD-based vs. DVARS-based thresholding

A consistent finding in the present study was that different frame censoring outcomes are obtained depending on whether FD or DVARS is used for thresholding. This effect is most striking in the maximum t-values observed in the individual studies (**Figure 3**). Systematically varying the FD and DVARS threshold values resulted in dissimilar or even contrary effects, with improvements observed in one metric often contrasting with worsening performance in the other. Although perhaps unexpected at first glance, this result reflects the nature of the two parameters and how censored frames are identified.

While FD is a direct quantification of estimated head motion, DVARS is potentially affected by any process that changes image intensity between frames. This includes not only head motion, but also both neural and non-neural influences such as arousal (Gu et al. 2020), respiration (Power et al. 2018), and cerebrospinal fluid flow (Fultz et al. 2019). As a result, even though FD and DVARS are strongly correlated, they are not identical, and this disparity is responsible for the observed differences in FD and DVARS performance. Even if the number of censored frames is equivalent (cf. **Figure 2d**), a different collection of frames is targeted by each parameter at a given threshold. The relationship between FD-based and DVARS-based thresholding can be conveniently demonstrated by considering the scatterplot of FD versus DVARS in **Figure 2b**. FD thresholding can be viewed in this plot as a vertical line moving right-to-left as the threshold is made more stringent. On the other hand, DVARS thresholding corresponds to a horizontal line moving top-to-bottom. Although there is a general overlap in the frames that violate both thresholds, the collections are not identical. Because the relation between the two parameters differs in each dataset (see **Supplemental Figure S3**), different trends in FD- and DVARS-based thresholded performance emerge.

#### Patterns of results across datasets

The similarity analysis of group-level maps (**Figure 5**) exhibits several notable features. First, untrained ICA had relatively low overlap with other motion correction strategies in most (but not all) datasets. Despite the frequently lower Dice scores, we did not see results for untrained ICA that were substantially mismatched with the other results. A review of the data reveals that the performance of untrained ICA seemed to result from less-extensive activation compared to group-level maps obtained using the other motion correction approaches. Stated differently, the untrained ICA activation maps were not “incorrect”; they were simply more focal (and thus overlapped less with other approaches).

Second, RP6 and RP24 produced a lower Dice overlap in many datasets. As these techniques are based on the use of continuous regressors, they represent an algorithmically distinct approach compared to temporally compact (wavelet) or discrete regressors (frame censoring). This effect can also be seen in the results of robust weighted least squares, which in some datasets (e.g., ds001497 and ds001534) produce the only notable Dice difference. As such, a tempting takeaway is that the motion correction strategies based on continuous regressors form a performance family. However, when all performance metrics are considered collectively, the distinction between approaches becomes less clear.

Finally, some of the overlap performance appears to be related to data quality. For example, ds001748 and ds002382 explored identical tasks across multiple samples of approximately equal size. Both datasets included a high-motion group (the children group in ds0001748 and the older adults in ds002382 — see **Table 1**) and it is these Dice matrices that exhibit the greatest variability within the group. Conversely, the Dice matrices for the ds001748 adult and teen subject pools and the young adults in ds002382 are relatively uniform. This suggests that the choice of a motion correction strategy may be less important when working with a subject pool exhibiting only minor motion, at least when considering the spatial distribution of group level activation.

These qualitative differences suggest Dice overlap might offer a means of categorizing the datasets and in so doing might provide a guideline for the selection of a motion correction strategy. A five-group categorization of the datasets can be proposed based simply on their appearance in **Figure 5**: 1) ds000102 and ds000114 (line bisection), 2) ds000114 (motor) and ds002382 (older adults), 3) ds000228 (adults), ds001748 (adults) and ds002382 (younger adults), 4) ds001748 (children), and 5) all remaining datasets. Yet, the quantitative results of our RDM-informed multidimensional scaling (**Figure 6** and **Supplementary Figure S4**) do not support this organization. Our goal was to identify common features of datasets using the overall pattern of motion correction results, which we operationalized using Dice overlap. However, this was not the case: MDS was unable to reduce the dimensionality of this data in a way that supplied meaningful information, and studies grouped together even using the informal visual organization described above differ in subject pools, task type, and other characteristics. Like the univariate metrics considered here, our multivariate analysis failed to clearly identify characteristics that might be used to identify an optimal motion correction strategy. It could be that a similar approach, but with hundreds of data sets, would be able to identify systematic differences in how different motion correction strategies worked on different types of data, which may be a promising direction for future work.

### Other considerations

We have focused on retrospective correction—that is, strategies for dealing with motion in existing data. A complementary approach would be to reduce head motion during acquisition. Protocols have been developed to do so, including movie viewing (Greene et al. 2018), custom head molds (Power et al. 2019), and providing feedback to participants (Dosenbach et al. 2017; Krause et al. 2019). However, these have not yet been widely adopted, nor are all compatible with task-based fMRI. With increasing awareness of the challenges caused by participant motion, perhaps greater interest in motion reduction (as opposed to motion correction) will follow.

A possibility which we did not explore is combining strategies, as is commonly done in resting state fMRI (e.g., frame censoring of outliers followed by including motion regressors from rigid body realignment). However, this expands an already unwieldy parameter space of possible analyses (Carp 2012; Poldrack et al. 2017; Botvinik-Nezer et al. 2020). The use of simulated data, where “ground truth” can be known, may also prove beneficial in understanding how motion correction strategy can affect the validity of our inferences.

## Conclusions

The present results do not identify unequivocal guidelines for selecting a motion correction strategy. Given the variability observed across datasets, analyzed using identical processing pipelines, exploring multiple strategies in a given dataset may be the best way of reducing motion artifacts. Although it may be possible to revisit this issue in future work, our present results suggest that—frustratingly—no single motion correction strategy will give optimal results in every instance, and that choices require considering both the nature of the specific data of interest and the most relevant outcome measure.

## Acknowledgments

This work was supported by grants R01 DC014281, R01 DC016594, R01 DC019507, and T32 EB014855 from the US National Institutes of Health. OpenNeuro is supported by NSF Grant OCI-1131441.

