## Supplemental materials for "A multi-dataset evaluation of frame censoring for motion correction in task-based fMRI"

### Supplemental Methods

#### Representational similarity analysis of group-level maps

Overlap of the thresholded group-level maps ($p < 0.001$, uncorrected) was quantified using a Dice coefficient computed for all pairs of denoising approaches. The result was a 16x16 matrix for each dataset in which the (i,j)-th entry is the Dice coefficient quantifying overlap of the group-level map obtained using the i-th denoising strategy with that of the j-th denoising strategy. The overlap summaries were then used to explore a multivariate characterization of the denoising strategies. To this end, all pairs of the 15 dice overlap matrices were correlated and the coefficients were collected into a 15x15 distance matrix (distance = 1-r). Multidimensional scaling (MDS; Matlab function *cmdscale*) was then applied to the distance matrix to obtain a low-dimensional approximation which was plotted and examined for clustering or other patterns.

#### Univariate statistical analysis

Maximum t values, mean ROI effect size, and Dice test-retest values for each denoising strategy were pooled across all datasets and analyzed using a repeated-measures analysis of variance (Matlab functions *fitrm* and *rmanova*). Maximum t, effect size and Dice test-retest score were normalized to the value obtained using no motion correction prior to pooling to account for variability across datasets. Pairwise differences were identified in post hoc testing using a Scheffe test (Matlab function *multcompare*).

An alternate evaluation of algorithm performance was explored by simply counting the number of datasets each approach gave the best result (i.e., the largest value) on a given metric. For the purpose of this analysis, FD and DVARS results were combined across all percent data loss categories, reducing the total number of categories to eight. Counts were then evaluated statistically using a binomial test (Zar, 2010). Under the null hypothesis that all denoising strategies perform equally well, exhibiting the best performance on seven or more datasets by any one strategy is significant at the 0.05 level.

### Supplemental Results

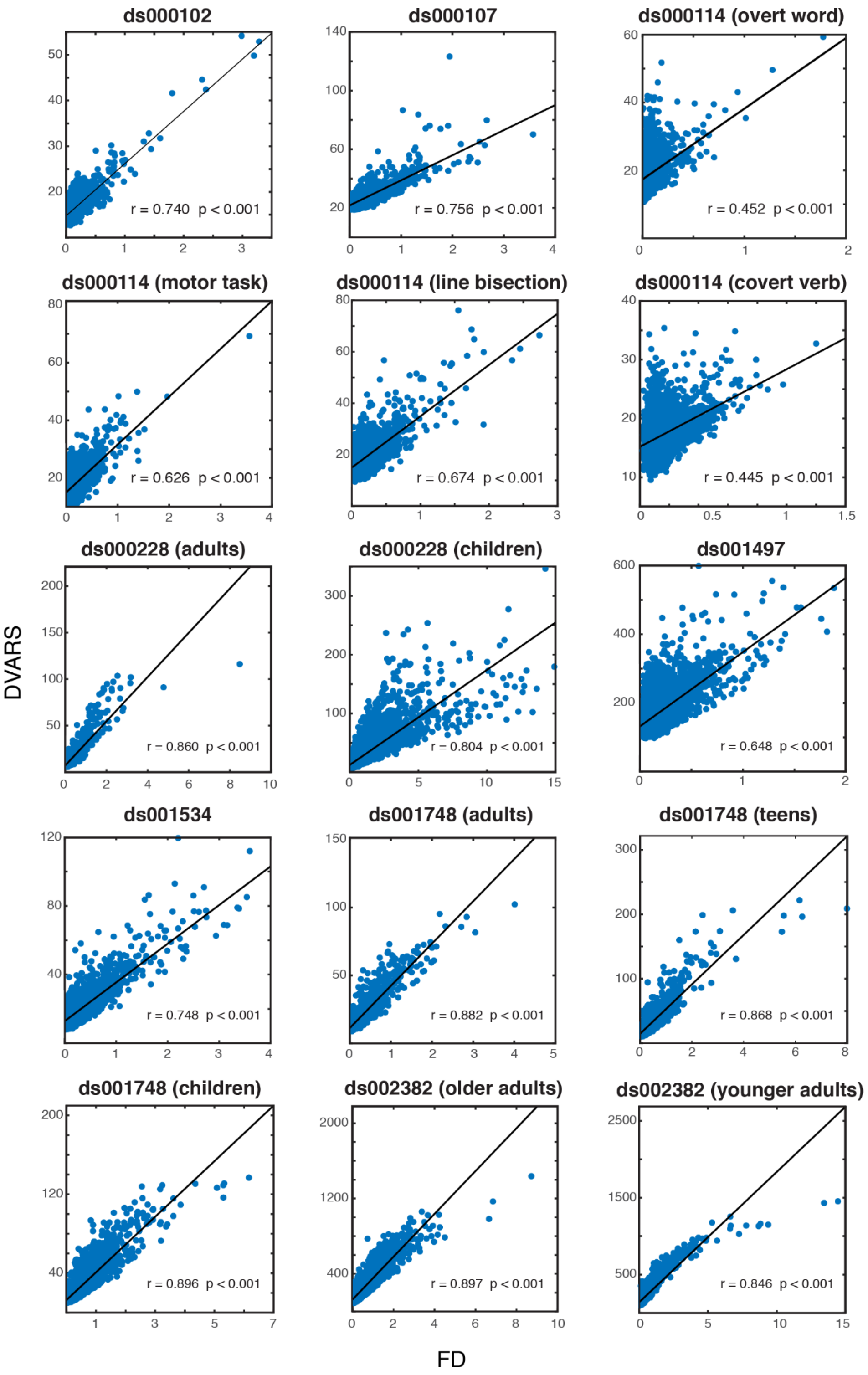


**Supplemental Figure S1.** Correlation of FD and DVARS for all datasets. Each point is the FD (x) and corresponding DVARS (y) for a single frame, combined across all sessions and subjects for a given dataset. Each plot includes a least-squares line fit to the data.

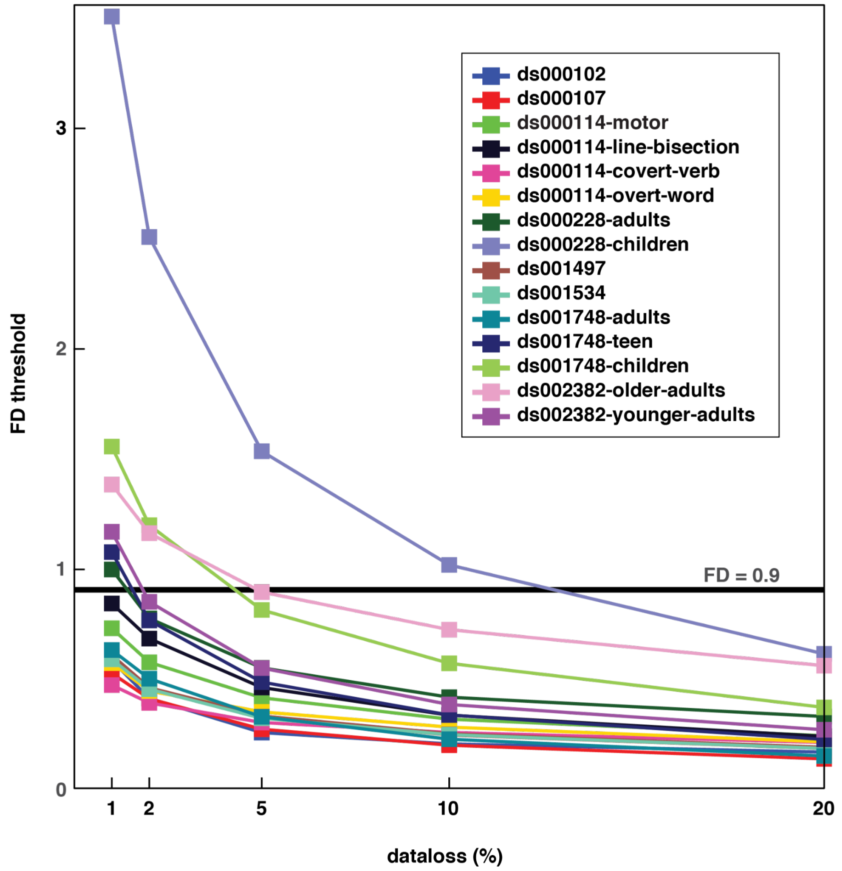


**Supplemental Figure S2. FD values resulting from targeted percent data loss.** The FD threshold value that resulted in 1, 2, 5, 10 or 20 percent data loss for a given dataset can be read from the y-axis. A horizontal line at FD = 0.9 is included to illustrate the data loss that would have occurred had an (arbitrary) fixed FD threshold of 0.9 been applied (given by the intersection or extrapolated intersection of the horizontal line with the graph of a dataset).

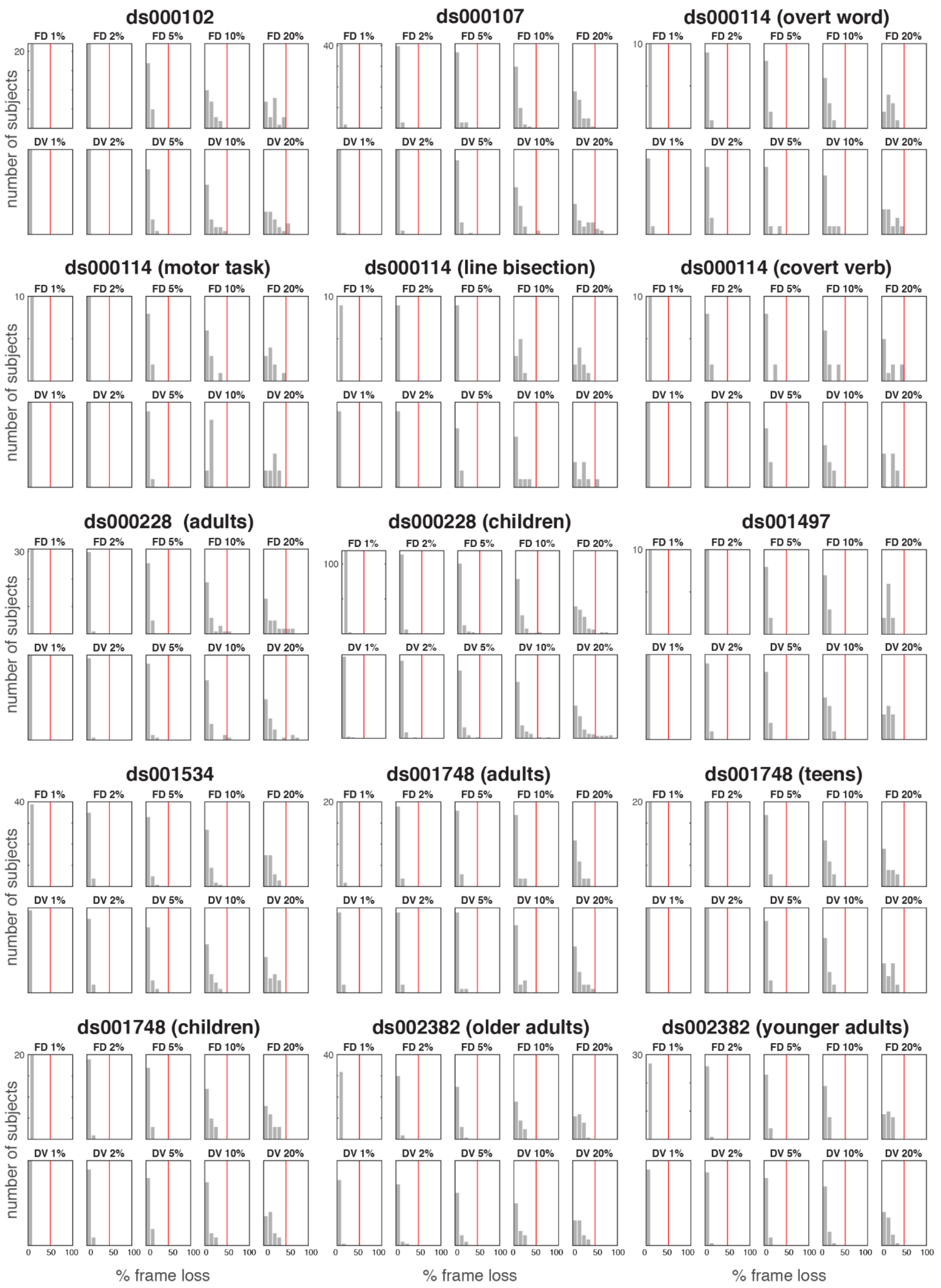


**Supplemental Figure S3:** Frameloss histograms for all 15 datasets and for all FD and DVARS thresholds. Red line indicates 50% frame loss for reference. See also Figure 2 in main text.

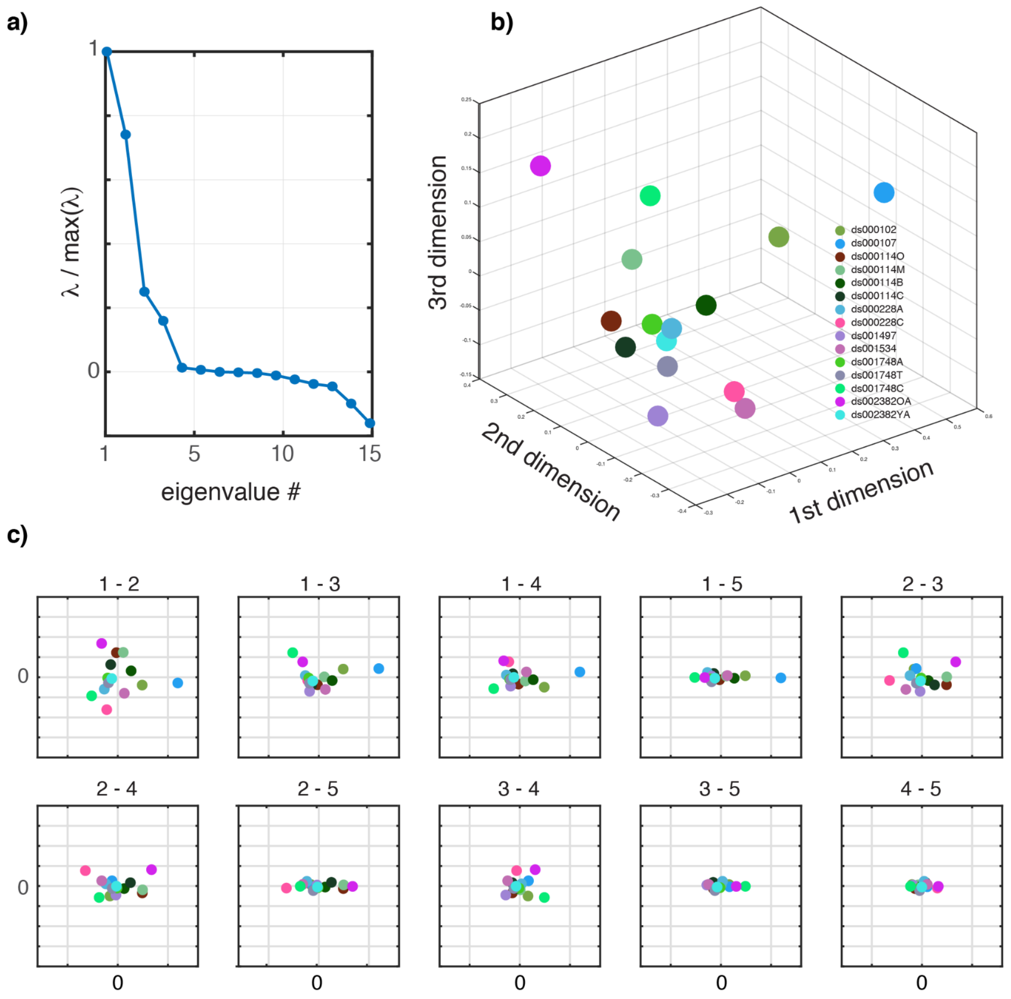


**Supplemental Figure S4.** Scree plot and additional MDS results. (a) Eigenvalue magnitude normalized to that of the largest eigenvalue returned by multidimensional scaling of the 15 Dice overlap matrices shown in Figure 4 in the main text. Falloff suggests the dimensionality of the distance data is about five. The presence of negative eigenvalues indicate the dataspace is non-Euclidean. (b) Plot of first three eigen-dimensions reveals no obvious pattern in the data. (c) Projections of all possible pairs of the first five eigen-dimensions. The dataset ds000107 is segregated in some of the plots, but no other organization is apparent.

**Supplemental Table 1. Some prior evaluations of motion correction in task-based fMRI.**

| Study | # Data sets | Task(s) | N | Motion correction approaches | Outcome measures |
|---|---|---|---|---|---|
| Current study | 8 | see Table 1 | 405 | RP6, RP24, rWLS, WDS, untrained ICA, FD- and DVAR-based frame censoring | whole brain and ROI maximum t, mean effect size, Dice |
| Diedrichsen and Shadmehr (2005) | 1 | hand controlled cursor targeting | 15 | robust weighted least squares | t-score, number of suprathreshold voxels |
| Hoffman et al. (2015) | 1 | auditory (SPM MoAE data contaminated with clinically relevant motion) | 1 | rigid body alignment in SPM, FSL, AFNI, or AIR | count of suprathreshold voxels, specious identification of motion |
| Huang et al. (2008) | 1 | reading aloud | 13 | linear interpolation over contaminated volumes | significant voxel counts |
| Johnstone et al. (2006) | 1 | go/no-go, N-back | 33 | rigid body registration with vs. without including six realignment parameters as covariates | whole-brain maximum t and cluster extent |
| Kay et al. (2013) | 11** | visual | 1-3 | custom (GLMDenoise), ICA | cross validation |
| Kochiyama et al. (2005) | 1 | finger tapping | 4 | ICA | ROC true vs false positive fraction |
| Lemmin et al. (2010) | 1 | arm motion | 7 | model-based motion estimation, RP6, rWLS | reduction in ventricle activation |
| Liao et al. (2006) | 1 | motor | 10 | custom ICA | standard deviation reduction in activated voxels |

| | | | | | |
|---|---|---|---|---|---|
| Mayer et al. (2019) | 2* | AX continuous performance task, multimodal attention | 110 | RP12, RP24, untrained ICA, trained ICA | percent change of true and false activation |
| Middlebrooks et al. (2017) | 1 | motor task, language task | 12 | realignment, DVARS scrubbing, trained ICA, ICA+scrubbing | increased z-scores in areas of expected activation |
| Oakes et al. (2005) | 1 | Go/No-go, N-back | 40 | rigid body registration in AFNI, AIR, BrainVoyager, FSL, or SPM | maximum t and cluster extent |
| Siegel et al. (2014) | 4* | string matching, rule switching, posner task | 88 | motion estimates as nuisance regressors vs. FD-thresholded frame censoring | change in beta estimate, error variance, or t-score |
| Tierney et al. (2016) | 1 | sentence comprehension and generation | 42 | custom algorithm (FIACH) compared with RP6, RP24, FD frame censoring, rWLS, tCompCorr | ROI restricted t-values and cluster extent |
| Tohka et al. (2008) | 2*** | tone counting, weather prediction | 32 | trained ICA | change in Z-scores |
| Wilke and Baldeweg (2019) | 3 | verb generation, hand motor task, language task | 84 | custom algorithm | t-value, # activated voxels, SNR |
| Xu et al. (2014) | 1 | overt speech | 18 | custom ICA | PET cross validation |

* same scanner, multiple cohorts
** 11 task variations run on a single cohort
*** a training and a test cohort on the same scanner/tasks

**Supplemental Table 2. Acquisition details for datasets analyzed**

| Dataset | Reference | Scanner | Field strength (T) | TR (s) | Voxel size (mm) |
|---|---|---|---|---|---|
| ds000102 | Kelly et al. (2008) | Siemens Allegra | 3 | 2 | 3x3x4 |
| ds000107 | Duncan et al. (2009) | Siemens Avanto | 1.5 | 3 | 3x3x3 |
| ds000114 | Gorgolewski et al. (2013) | GE Signa HDxt | 1.5 | 2.5 | 4x4x4 |
| ds000228 | Richardson et al. (2018) | Siemens Tim Trio | 3 | 2 | 3x3x3 |
| ds001497 | Lewis-Peacock and Postle (2008) | GE Signa VH/I | 3 | 2 | 3.75x3.75x4 |
| ds001534 | Courtney et al. (2018) | Philips Intera Achieva | 2.5 | 2.5 | 3x3x3 |
| ds001748 | Fynes-Clinton et al. (2019) | Siemens Magnetom Trio | 3 | 3 | 2.5x2.5x2.5 |
| ds002382 | Rogers et al. (2020) | Siemens Prisma | 3 | 3.07 | 2x2x2 |

**Supplemental Table 3. DVARS and FD values at percent frameloss**

| | FD | | | | | DVARS | | | | |
|---|---|---|---|---|---|---|---|---|---|---|
| **Dataset** | **1%** | **2%** | **5%** | **10%** | **20%** | **1%** | **2%** | **5%** | **10%** | **20%** |
| ds000102 | 22.1 | 20.2 | 18.5 | 17.5 | 16.9 | 0.58 | 0.41 | 0.25 | 0.20 | 0.16 |
| ds000107 | 32.3 | 29.5 | 26.8 | 25.3 | 24.2 | 0.52 | 0.41 | 0.27 | 0.20 | 0.13 |
| ds000114 bisection | 35.4 | 30.8 | 25.9 | 23.1 | 20.8 | 0.84 | 0.68 | 0.46 | 0.33 | 0.24 |
| ds000114 covert verb | 25.5 | 23.8 | 22.1 | 20.6 | 19.0 | 0.47 | 0.39 | 0.30 | 0.25 | 0.21 |
| ds000114 lips motor | 31.1 | 28.1 | 24.3 | 22.1 | 20.1 | 0.73 | 0.57 | 0.41 | 0.31 | 0.24 |
| ds000114 overt word | 33.7 | 31.8 | 28.6 | 26.3 | 23.7 | 0.57 | 0.44 | 0.35 | 0.28 | 0.21 |
| ds000228 adults | 30.1 | 25.3 | 20.9 | 17.4 | 15.1 | 0.99 | 0.77 | 0.55 | 0.41 | 0.33 |
| ds000228 children | 72.3 | 59.1 | 42.1 | 31.6 | 23.5 | 3.50 | 2.50 | 1.53 | 1.01 | 0.61 |
| ds001497 | 284.6 | 251.3 | 219.4 | 198.3 | 179.6 | 0.60 | 0.45 | 0.33 | 0.24 | 0.18 |
| ds001534 | 28.2 | 24.1 | 20.7 | 18.8 | 17.1 | 0.59 | 0.45 | 0.32 | 0.24 | 0.18 |
| ds001748 adults | 31.0 | 27.4 | 22.0 | 19.2 | 16.4 | 0.63 | 0.50 | 0.32 | 0.22 | 0.15 |
| ds001748 children | 58.0 | 50.5 | 38.4 | 30.0 | 23.5 | 1.55 | 1.19 | 0.81 | 0.57 | 0.37 |
| ds001748 teen | 58.7 | 50.1 | 35.9 | 28.6 | 23.7 | 1.07 | 0.76 | 0.48 | 0.33 | 0.22 |
| ds002382 older adult | 463.4 | 410.1 | 343.6 | 300.8 | 258.0 | 1.38 | 1.16 | 0.89 | 0.72 | 0.56 |
| ds002382 young adult | 382.0 | 324.3 | 258.8 | 222.1 | 193.2 | 1.16 | 0.85 | 0.55 | 0.38 | 0.27 |

**Additional Dataset Details**

Details of datasets used in the study are summarized below, including challenges or irregularities we encountered during analysis.

*Accession Number: ds000102*
Publication: Kelly et al. (2008)
Task: Slow event-related Eriksen flanker
Task details: Participants used one of two buttons to indicate the direction of a central arrow in an array of five arrows. In congruent trials, the flanking arrows pointed in the same direction as the central arrow; in more demanding incongruent trials the flanking arrows pointed in the opposite direction.
Acquisition: Siemens Allegra 3.0 T (TR = 2000 ms; TE = 30 ms; flip angle = 80, 40 slices, matrix = 64x64; FOV = 192 mm; acquisition voxel size = 3 × 3 × 4 mm).
Number of subjects: 26; Age: 22-50 years (mean 32)
Contrast evaluated: incongruent correct > congruent correct

ROI: Neuroquery search term "incongruent task" and thresholded Z > 3
Notes: Functional data includes a pronounced periodic artifact that appears unrelated to motion.

*Accession Number: ds000107*
Publication: Duncan et al. (2009)
Task: One-back
Task details: A one-back task was used with four categories of visual stimuli: written words, pictures of common objects, scrambled pictures of the same objects, and consonant letter strings. Subjects were instructed to press a button if the stimulus was identical to the preceding stimulus (12.5% of the stimuli were targets). Each block consisted of 16 trials from a single category presented one every second. A trial began with a 650 ms fixation cross, followed by the stimulus for 350 ms.
Acquisition: Siemens Avanto 1.5 T. The functional data were acquired with a gradient-echo EPI sequence (TR = 3000 ms; TE = 50 ms; FOV = 192 × 192; matrix = 64 × 64, voxel size = 3 × 3 × 3 mm).
Number of subjects: 45 (23 Male); Age: 19-38 years (mean 25)
Contrast evaluated: Words > 0
ROI: 5 mm sphere at [-42 -62 -16] (left ventral occipital-temporal cortex). Coordinates are from Table 2 of Duncan et al. (2009).
Notes: Data from six subjects were excluded: two because of corrupt data, three due to data modeling errors that could not be corrected, and one due to an incompatible contrast definition. Removed spaces from event names in BIDS .tsv files.

*Accession Number: ds000114*
Publication: Gorgolewski et al. (2013)
Tasks: i) lip movement, ii) covert verb generation, iii) overt word generation, iv) line bisection. All tasks used the same subjects.
Task details: Lip movement: Lip poaching (15 s) interleaved with fixation at a cross (15 s). Covert verb generation: Subjects instructed to think of a verb following presentation of a random noun for 1 s. Overt word generation: Repeat words aloud presented via headphones; 30 s task / 30 s rest repeated six times. Line bisection: Judge by button press if a horizontal line was bisected exactly in the middle (landmark) or if a horizontal line was crossed or not crossed (detection). Randomized presentation of six correct and four incorrect lines (525 ms presentation / 1100 ms response) in eight blocks.
Acquisition: GE Signa HDxt 1.5 T scanner with an 8-channel phased-array head coil. FOV = 256 × 256 mm, voxel size = 4 x 4 x 4 mm, slice thickness 4 mm, 30 slices per volume, interleaved slices order, acquisition matrix 64 × 64, flip angle = 90, TE = 50 ms, TR = 2.5 s, except for overt word repetition in which sparse sampling was used (TR = 5 s, “real TR”—which we assumed meant “acquisition time/TA” = 2.5 s). Subjects were scanned twice, either two or three days apart.
Number of subjects: 10 (4 Male); age: 50-58 years (median 52.5).
Contrast evaluated: i) Motor task: lip > hand+foot, ii) Covert verb: task > 0, iii) Overt word: task > 0, iv) Line bisection: landmark > detection
ROI: i) 5 mm spheres in bilateral motor cortex ([-56,-6,26] and [62,0,28]). Coordinates taken from (Pulvermüller et al., 2006), ii) Covert verb: Broca's Area (BA 44 + BA 45 (left hemisphere

only) from Anatomy Toolbox), iii) Overt Word: same as lip motor task, iv) Line bisection: lateral visual cortex (Anatomy Toolbox hoc3v+doc4v+hoc4lp).
Notes: The motor data included finger and foot tapping tasks that were not used because a mixture of left- and right-handed activation precluded straightforward second-level modeling. Subject 10 was excluded from the line bisection analysis as first level maps suggest the subject misunderstood task instructions (activation is left/right reversed).

*Accession Number: ds000228*
Publication: Richardson et al. (2018)
Task: Film viewing
Task details: Subjects viewed a 5.6-minute animated film with scenes classified as presenting either "pain" or "theory of mind" events.
Acquisition: 3T Siemens Tim Trio using a standard Siemens 32-channel head coil. T1-weighted structural images were collected in 176 interleaved sagittal slices with 1 mm isotropic voxels (GRAPPA parallel imaging, acceleration factor of 3; adult coil: FOV: 256 mm; kid coils: FOV: 192 mm). Functional data were collected with a gradient-echo EPI sequence in 32 interleaved near-axial slices aligned with the anterior/posterior commissure, and covering the whole brain (EPI factor: 64; TR: 2 s, TE: 30 ms, flip angle: 90). Voxel size: Adults 3.13 × 3.13 × 3.13 mm; children either 3 × 3 × 3 mm or 3.13 × 3.13 × 3.13 mm.
Number of subjects: Adults: 33 (20 female); age: 18–39 years (mean: 24.8). Children: 123 (64 female); age: 3.5–12 years; mean: 6.7).
Contrast evaluated: pain > theory of mind
ROI: Regions listed in Supplementary Table 2 of Richardson et al. (2018), modeled as a collection of 5 mm spheres.
Notes: There were insufficient sessions in this data for test-retest evaluation. Event files were missing from the original OpenNeuro listing and were added manually using information provided in the description. The event timing provided was converted to seconds from scans using the TR information.

*Accession Number: ds001497*
Publication: Lewis-Peacock and Postle (2008)
Task: Stimulus judgment / memory
Task details: Subjects viewed a total of 90 stimuli drawn from three categories: 30 famous people, 30 famous locations, and 30 common objects. They indicated (on a four-point Likert scale, using a stimulus-response box) how much they liked the celebrity, how much they would like to visit the location, or how often they encountered the object in everyday life.
Acquisition: GE Signa VH/I 3T scanner. T1 (30 axial slices, 0.9375 × 0.9375 × 4 mm).
Functional images: gradient-echo echo-planar (TR = 2000 ms; TE = 50 ms; 64 x 64 matrix coplanar with the T1 acquisition, voxel size = 3.75 × 3.75 × 4 mm).
Number of subjects: 10 (7 male); age: 19-32 years.
Contrast evaluated: Face > 0
ROI: Bilateral fusiform face area (Neuroquery search term "FFA" and thresholded Z > 3)
Notes: There were a total of six sessions in the data which were split into even and odd sessions for test-retest evaluation. The data on OpenNeuro is only the "LTM" portion of the experiment. Data from a working memory task described in the associated publication is not

included.

*Accession Number: ds001534*
Publication: Courtney et al. (2018)
Task: Food images paired with textural calorie content
Task details: Participants first viewed images of food paired with an accompanying image number ("foodimage"), and subsequently viewed these same food images paired with the corresponding calorie information ("calorieimage"). The presentation sequence of food images and jittered fixation trials were pseudo-randomized.
Acquisition: Philips Intera Achieva scanner. Anatomical images were acquired using gradient-echo sequence (TR = 9.9 ms; TE = 4.6 ms; flip angle = 8; 1x1x1 mm voxels). Functional images were collected using T2* fast field echo (TR = 2.5 seconds, TE = 35 ms, flip angle = 90, voxel size = 3 × 3 × 3 mm).
Number of subjects: 50 (50 M); age: 18-22 years (mean 19.7).
Contrast evaluated: labeled image > not-labeled
ROI: Bilateral inferior parietal cortex (Anatomy Toolbox IPC_PF + IPC_PFcm + IPC_PFm + IPC_PFop + IPC_PFt + PIC_PGa + IPC + PGp).
Notes: Functional images in this data were scaled by 0.02 prior to processing.

*Accession Number: ds001748*
Publication: Fynes-Clinton et al. (2019)
Task: Memory retrieval including autobiographical, episodic, or semantic conditions
Task details: One of 25 images of everyday life events were presented for 4 s, followed by a retrieval cue screen for 8 seconds during which participants retrieve different long-term memories. The type of memory retrieval was manipulated by adjusting the response screen to cue the retrieval of either personal experience (AM), general knowledge and factual information (SM), or questions about the content of the cue images (EM).
Acquisition: 3T Siemens scanner equipped with a 32-channel head coil. Structural: 176 slices sagittal; 1 mm isotropic volume; TR = 4000 ms; TE = 2.89 ms; FOV = 256 mm. Functional: T2*-weighted echo-planar image pulse sequence (45 slices, 2.5 mm slice thickness; voxel size = 2.5 × 2.5 × 2.5 mm, TR = 3000 ms; TE = 30 ms; FOV = 190 mm; flip angle = 90).
Number of subjects: 62 (32M); age: 10-35 years (see Notes).
Contrast evaluated: task > control
ROI: Inferior frontal gyrus (Neuroquery search term "IFG" thresholded Z > 3)
Notes: Data comprised three cohorts: children (10-12; n=21), adolescents (14-16; n=20) and young adults (20-35; n=22) that were analyzed separately. There was insufficient data for test-retest evaluation. The tsv file for child-20 contains a typo with "semantic" mislabeled as "semanti" and autobio.tsv is empty for child-13. These subjects were excluded.

*Accession Number: ds002382*
Publication: Rogers et al. (2020)
Task: Speech comprehension in noise
Task details: Subjects were presented auditory stimuli via MR-compatible headphones consisting of words (monosyllabic consonant-vowel-consonant), silence, and noise (single-channel noise vocoded words) in two sessions of passive listening and two session of word

repetition in which participants were asked to repeat aloud the presented word. Responses in the repeat condition were recorded and scored as either correct or incorrect.

Acquisition: MRI data were acquired using a Siemens Prisma scanner (Siemens Medical Systems) at 3 T equipped with a 32-channel head coil. Scan sequences began with a T1-weighted structural volume using an MPRAGE sequence (TR = 2.4 s, TE = 2.2 ms, flip angle = 8, 300 × 320 matrix, voxel size = 0.8 mm isotropic). Functional images were acquired using a multiband echo planar imaging sequence (TR = 3.07 s, TA = 0.770 s, TE = 37 ms, flip angle = 37, voxel size = 2 × 2 × 2 mm, multiband factor = 8).

Number of subjects: Young adults: n = 29 (19 female); age: 19–30 years (mean = 23.8). Older adults: n = 32 (17 female); age 65–81 years (mean = 71.0).

Contrast evaluated: repeat word > noise

ROI: 5 mm spheres in bilateral motor cortex ([-56,-6,26] and [62,0,28]). Coordinates taken from Pulvermüller et al. (2006)

Notes: Young adults and older adults were analyzed separately.
